# Nanoparticle delivery of innate immune agonists combines with senescence-inducing agents to mediate T cell control of pancreatic cancer

**DOI:** 10.1101/2023.09.18.558307

**Authors:** Loretah Chibaya, Christina F. Lusi, Kelly D. DeMarco, Griffin I. Kane, Meghan L. Brassil, Chaitanya N. Parikh, Katherine C. Murphy, Junhui Li, Tiana E. Naylor, Julia Cerrutti, Jessica Peura, Jason R. Pitarresi, Lihua Julie Zhu, Katherine A. Fitzgerald, Prabhani U. Atukorale, Marcus Ruscetti

## Abstract

Pancreatic ductal adenocarcinoma has quickly risen to become the 3^rd^ leading cause of cancer-related death. This is in part due to its fibrotic tumor microenvironment (TME) that contributes to poor vascularization and immune infiltration and subsequent chemo- and immunotherapy failure. Here we investigated an innovative immunotherapy approach combining local delivery of STING and TLR4 innate immune agonists *via* lipid-based nanoparticles (NPs) co-encapsulation with senescence-inducing RAS-targeted therapies that can remodel the immune suppressive PDAC TME through the senescence-associated secretory phenotype. Treatment of transplanted and autochthonous PDAC mouse models with these regimens led to enhanced uptake of NPs by multiple cell types in the PDAC TME, induction of type I interferon and other pro-inflammatory signaling, increased antigen presentation by tumor cells and antigen presenting cells, and subsequent activation of both innate and adaptive immune responses. This two-pronged approach produced potent T cell-driven and Type I interferon-dependent tumor regressions and long-term survival in preclinical PDAC models. STING and TLR4-mediated Type I interferon signaling were also associated with enhanced NK and CD8^+^ T cell immunity in human PDAC. Thus, combining localized immune agonist delivery with systemic tumor-targeted therapy can synergize to orchestrate a coordinated innate and adaptive immune assault to overcome immune suppression and activate durable anti-tumor T cell responses against PDAC.

**SUMMARY:** Combining senescence-inducing MEK and CDK4/6 inhibitors with nanoparticle delivery of STING and TLR4 agonists leads to interferon-driven and cytotoxic T cell-mediated PDAC control.

## INTRODUCTION

PDAC is a devastating disease with a dismal 5-year survival rate of 11% (*1*), largely due to the physiological makeup of the TME that promotes tumor advancement and limits effective treatment options. Characterized by a hallmark desmoplastic stroma, poor vascularization, and immunosuppression, the PDAC TME hinders effective drug delivery, drives chemo-resistance, and blocks the activation and infiltration of cytotoxic immune cells (*2*). Though immune checkpoint blockade (ICB) therapies targeting inhibitory checkpoints such as PD-1 and CTLA-4 on T cells have demonstrated durable responses in some cancer types, they have not shown efficacy in the immune suppressed PDAC TME that is devoid of CD8^+^ cytotoxic T lymphocytes (CTLs) and Natural Killer (NK) cells as well as antigen-presenting cells (APCs) such as dendritic cells (DCs) necessary to sustain anti-tumor T cell immunity (*3–6*). In addition, pancreatic tumor cells themselves have poor immunogenicity and antigenicity mediated in part by suppression of interferon signaling by oncogenic KRAS driver mutations (*7*). Thus, combinatorial approaches targeting the multi-faceted immune suppressive network of the PDAC TME are warranted to effectively orchestrate CTL-mediated anti-tumor immunity.

We and others have demonstrated that RAS-targeted therapies not only increase antigen presentation through upregulation of major histocompatibility complex (MHC) Class I (MHC-I) molecules on tumor cells, but can also induce cellular senescence and a subsequent senescence-associated secretory phenotype (SASP) including angiogenic and inflammatory factors that can remodel immune suppressive TMEs in dynamic ways (*8–14*). In KRAS mutant lung adenocarcinoma (LUAD) models, treatment with combinations of the MEK inhibitor trametinib (T) and CDK4/6 inhibitor palbociclib (P) that target downstream KRAS signaling induce a pro-inflammatory SASP leading to NK cell-mediated lung tumor regressions (*12*). In contrast, treatment with the same T/P regimens in KRAS mutant PDAC models induced a pro-angiogenic SASP that promoted vascular remodeling and endothelial activation leading to increased chemotherapy delivery, CTL trafficking, and anti-PD-1 ICB efficacy (*13*). These organ-specific differences in immune responses appear to be dictated by the resident microenvironment. In particular, we recently uncovered that myofibroblasts prevalent in the PDAC TME contribute to suppression of pro-inflammatory SASP factors and effective NK and CD8^+^ T cell immunity following T/P treatment (*15*). Notably, targeting the mechanisms of TME-driven immune suppression led to reactivation of interferon regulatory factor (IRF) expression and downstream interferon signaling that are normally induced in LUAD but repressed in PDAC, suggesting that approaches to engage IFN signaling could be a means to activate CTL activity in the PDAC.

The Stimulator of Interferon Genes (STING) pathway is a major regulator of type I interferon production and has emerged as an important innate immune pathway that can enhance anti-tumor NK and T cell immunity (*16*). Upon binding 2′-3′-cyclic-GMP-AMP (cGAMP), a second messenger produced by cyclic GMP–AMP synthase (cGAS) following recognition of cytoplasmic double-stranded DNA, STING stimulates downstream activation of IRF3 and NF-κB transcriptional activity to drive type I interferon (e.g. IFNβ) and other pro-inflammatory cytokines and chemokines, including those associated with the SASP (*17–20*). Administration of cGAMP and other synthetic STING agonists have been shown to stimulate antigen-presenting DCs, reprogram immune suppressive macrophages, reduce inhibitory regulatory T cell (Treg) numbers, and increase CD8^+^ T cell activation, leading to anti-tumor effects in preclinical PDAC models (*21–23*). Though promising, the clinical development of STING agonists as potential immunotherapies has been constrained by (a) their unfavorable pharmacokinetics and poor bioavailability due to limited cellular uptake and half-life in circulation, (b) adverse toxicities and immune suppressive effects associated with systemic administration, and (c) the inaccessibility of some tumor sites, including the pancreas, to intratumoral administration (*24–26*).

To overcome these limitations, we have designed lipid-based nanoparticles (NPs) that allow for systemic delivery of payloads of STING agonists that are preferentially deposited and taken up by APCs within the “leaky” perivascular region of tumors because of their “stealth” surface coating and small size (*27*). Taking advantage of the fact that nanoparticles can deliver multiple immune-stimulating agonists as cargos, NPs were loaded with not only the STING agonist cyclic di-guanosine monophosphate (cdGMP), but also the Toll-like receptor 4 (TLR4) agonist monophosphoryl lipid A (MPLA) that can additionally stimulate Type I interferon responses (*28, 29*). We have shown in melanoma and triple-negative breast cancer models that systemic co-delivery of STING and TLR4 agonists in NPs (hereafter immuno-NPs) drives their access to and uptake into the TME of even poorly vascularized tumors and leads to synergistic and robust IFNβ production as compared to NPs carrying either single agonist alone (*27, 30–32*). Immuno-NP administration in these models resulted in potent activation of innate (APCs, NK cells) and adaptive (CTL) immune responses, reduced tumor growth, and prolonged survival that could not be achieved with free agonist delivery or even anti-PD1 ICB (*27, 30–32*).

We hypothesized that combining T/P and immuno-NP therapy would orchestrate a coordinated remodeling of immune suppressive networks within immune cells, tumor cells, and the vasculature in the PDAC TME to produce durable anti-tumor T cell responses. Here, using syngeneic transplant and autochthonous PDAC mouse models, we found that combined treatment led to enhanced immuno-NP uptake in multiple cell types in the TME, synergistic activation of both type I interferons and SASP-associated cytokines and chemokines, and upregulated antigen presentation on tumor cells and APCs that culminated in sustained interferon alpha and beta receptor subunit 1 (IFNAR)-dependent and CD8^+^ T cell-mediated anti-tumor immune responses against PDAC.

## RESULTS

### NP encapsulation facilitates effective delivery of STING and TLR4 agonists to multiple cell types in PDAC TME

We first set out to assess whether STING and TLR4 agonists could be locally delivered to the PDAC TME through systemic intravenous administration following their co-encapsulation in lipid-based nanoparticles (immuno-NPs). Specifically, immuno-NPs were engineered with a ∼40-nm diameter and “stealth” poly(ethylene) glycol (PEG) surface to enable safe and effective delivery in the systemic blood circulation. These lipid-based materials supported the co-loading of the hydrophilic STING agonist, cdGMP, in the aqueous core and the hydrophobic TLR4 agonist, MPLA, in the lipid bilayer shell (Fig. 1, A to C). A lipid fluorescent Di tracer was also incorporated into the bilayer to track and assess NP biodistribution *in vivo*. Fluorescent NPs were then administered systemically into the bloodstream by tail vein injection to either tumor-bearing (a) C57BL/6 mice orthotopically transplanted with *KPC* PDAC cell lines engineered with a luciferase-GFP reporter to track them *in vivo* or (b) *P48-Cre*;*Kras^LSL-G12D/wt^*;*Trp53^fl/wt^*(*KPC*) genetically engineered mouse models (GEMMs) that spontaneously develop PDAC.

**Fig. 1.**
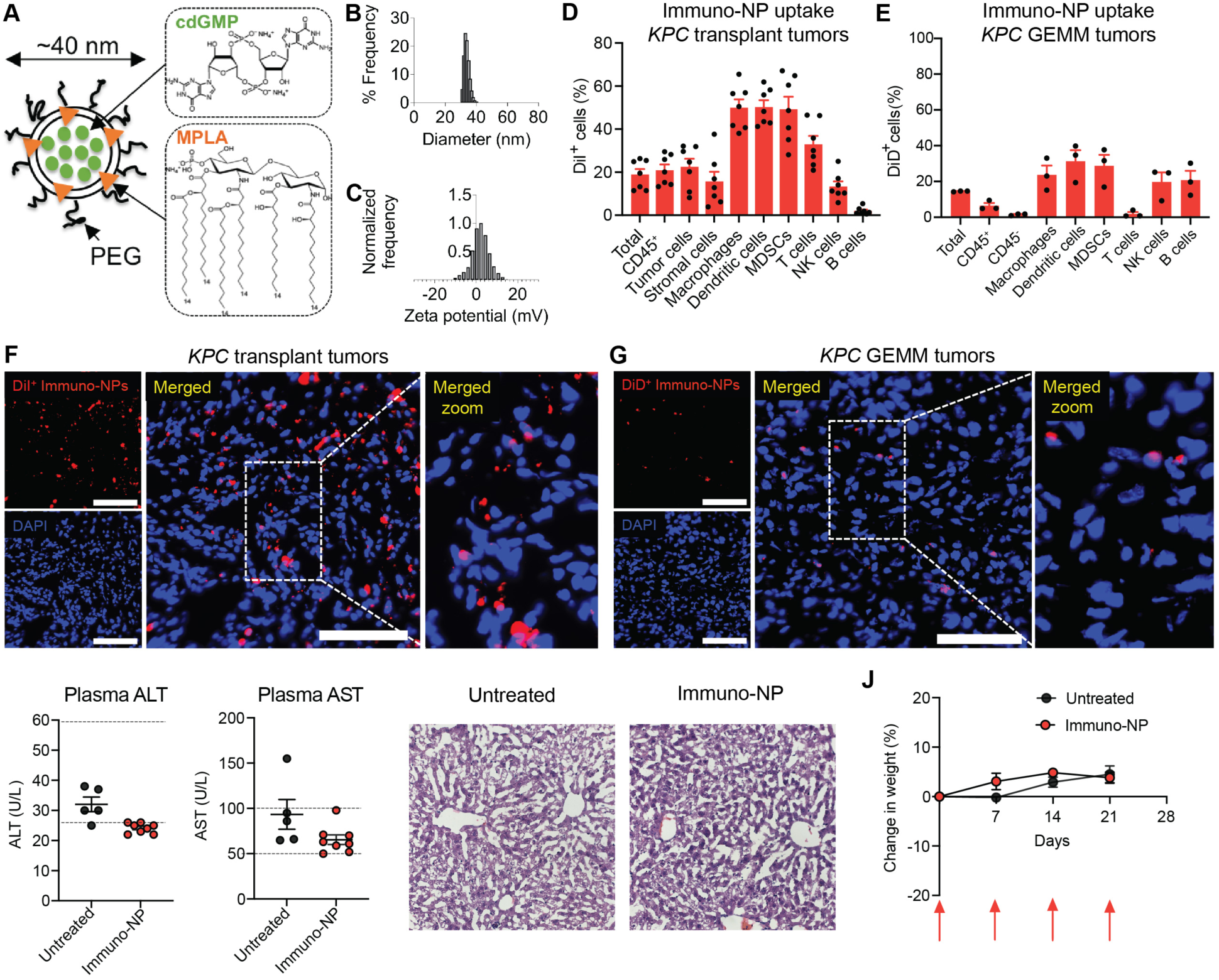
Systemic administration of NPs can deliver cargo locally to multiple cell types in PDAC TME with minimal toxicity. (**A**) Schematic representation of immuno-NP design. (**B**) NP hydrodynamic size as assessed by dynamic light scattering (DLS). (**C**) Measurement of NP surface charge as assessed by zeta potential. (**D**) *KPC1* PDAC tumor cells expressing luciferase-GFP were injected orthotopically into the pancreas of 8-12 week old C57BL/6 female mice. Following tumor formation, mice received a single dose of fluorescently labeled immuno-NPs by intravenous (i.v.) injection. Flow cytometry analysis of DiI^+^ NP uptake in indicated cell types 48hrs later is shown (*n* = 7 mice per group). Tumor cells were defined as GFP^+^, and stromal cells as CD45^-^GFP^-^. (**E**) PDAC-bearing *KPC* GEMM mice were i.v. injected with a single dose of fluorescently labeled immuno-NPs. Flow cytometry analysis of DiD-labeled NP uptake in different cell types 48hrs later is shown (*n* = 3 mice per group). (**F**) Representative immunofluorescence (IF) staining of *KPC1* orthotopic transplant PDAC tumors for expression of DiI-labeled immuno-NPs. Scale bars, 100 µm. (**G**) Representative immunofluorescence (IF) staining of *KPC* GEMM PDAC tumors for expression of DiD-labeled immuno-NPs. Scale bars, 100 µm. (**H**) Plasma AST and ALT levels in Wild-type (WT) C57BL/6 mice either untreated or treated with immuno-NPs weekly for 3 weeks (*n* = 5 to 8 mice per group). Dotted lines indicate established range for normal AST and ALT levels. (**I**) Representative Hematoxylin and eosin (H&E) staining of livers from WT C57BL/6 mice treated as in (H). (**J**) Change in tumor weight of WT C57BL/6 mice treated as in (H) (*n* = 5 to 8 mice per group). Arrows indicate when immuno-NPs were administered. Error bars, mean ± SEM.

NPs could be detected in the PDAC TME in both transplant and autochthonous PDAC models 48 hours post-injection. In *KPC* transplant mice, NPs were taken up by ∼20% of live cells in the PDAC TME and preferentially, as predicted, by myeloid cells such as macrophages, DCs, and myeloid-derived suppressor cells (MDSCs) that are enriched in perivascular regions (Fig. 1, D and F, and fig. S1). Still, other immune cell populations, as well as tumor cells and CD45^-^ stromal cells, also stained positive for the fluorophore-labeled NPs (Fig. 1D). NPs were also successfully delivered to myeloid cells and other immune populations in the PDAC TME of densely fibrotic autochthonous *KPC* GEMMs that are notoriously difficult for drugs to penetrate (Fig. 1, E and G). Importantly, NP-mediated delivery of these innate immune agonists led to no observable liver toxicity (Fig. 1, H and I) or weight loss (Fig. 1J), even after repeated weekly dosing of mice. Collectively, these results demonstrate that NPs can safely deliver immune stimulatory STING and TLR4 agonists to multiple cell types in the PDAC TME.

### Combinatorial immuno-NP and T/P treatment leads to synergistic induction of IFN and inflammatory signaling and antigen presentation in tumor cells and APCs

As we previously demonstrated that the MEK inhibitor trametinib and CDK4/6 inhibitor palbociclib (T/P) induce senescence and a pro-angiogenic SASP leading to vascular remodeling and increased drug uptake in PDAC lesions (*13*), we hypothesized that combining T/P with immuno-NP treatment would further enhance NP biodistribution. Indeed, pre-treatment with T/P for 12 days increased the uptake of immuno-NPs, as well as control unloaded empty NPs, by multiple tumor and immune cell types in the PDAC TME of *KPC* transplant mice (Fig. 2, A and B). Consequently, combined T/P and immuno-NP treatment significantly increased expression of not only downstream STING pathway components (*Tbk1*, *Irf3*) and the Type I interferon *Ifnb1*, but also pro-inflammatory SASP regulators (*p65*) and factors that we have previously shown to be repressed in the PDAC TME (*15*) compared to either treatment alone, including cytokines (*Il12, Il18*) and chemokines (*Ccl2, Ccl3, Cxcl10, Cx3cl1*) important for the activation and infiltration of cytotoxic NK and T lymphocytes (Fig. 2C). Co-immunofluorescence staining in both *KPC* transplant and GEMM PDAC lesions revealed the strongest induction of IFNβ following T/P and immuno-NP treatment not just in APCs such as DCs and macrophages, but also in tumor cells where it is not normally expressed (Fig. 2D and fig. S2A).

**Fig. 2.**
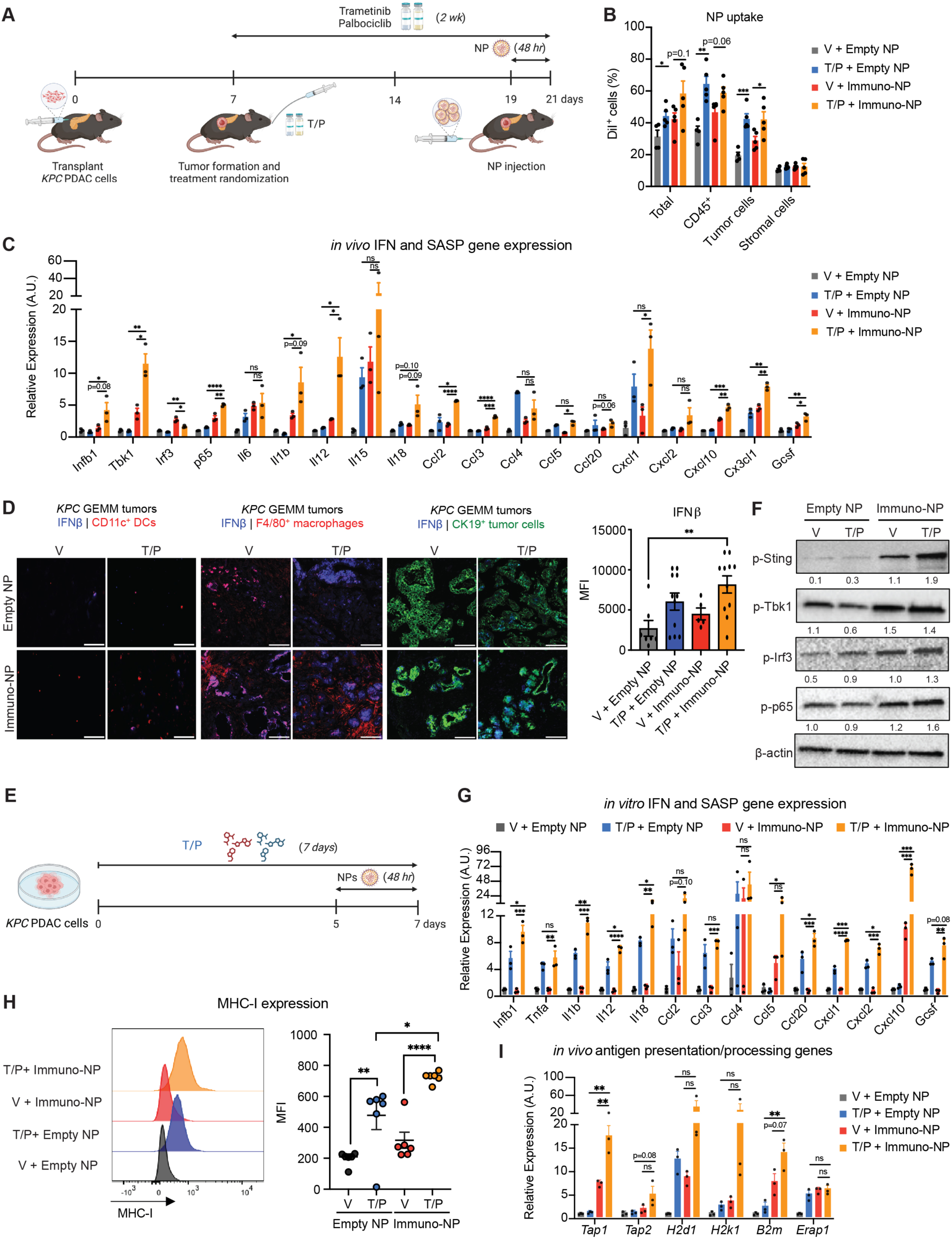
T/P pre-treatment enhances immuno-NP uptake, IFN and cytokine production, and antigen presentation in tumor cells and APCs. (**A**) Schematic of *KPC* orthotopic transplant model and 2-week treatment schedule. (**B)** Flow cytometry analysis of DiI-labeled NP uptake in indicated cellular compartments in *KPC1* transplant PDAC tumors from mice treated with vehicle or trametinib (1 mg/kg) and palbociclib (100 mg/kg) for 2 weeks and empty-or immuno-NPs for 48 hrs (*n* = 4 to 5 mice per group). Tumor cells were defined as GFP^+^, and stromal cells as CD45^-^ GFP^-^. (**C**) RT-qPCR analysis of IFN pathway and SASP gene expression in *KPC1* transplant PDAC tumors from mice treated as in (B) (*n* = 3 mice per group). A.U., arbitrary units. (**D**) Representative IF staining of *KPC* GEMM PDAC tumors from mice treated as in (B) for expression of IFNβ in DCs (CD11c^+^), macrophages (F4/80^+^), and tumor cells (CK19^+^). Quantification of mean fluorescent intensity (MFI) of total IFNβ expression in tissues is shown in last panel on right (*n* = 3 mice per group). Scale bars, 100 µm. (**E**) Schematic of *KPC1* cell line *in vitro* treatment schedule. (**F**) Immunoblots of *KPC1* PDAC cells treated *in vitro* with vehicle or trametinib (25 nM) and palbociclib (500 nM) for 1 week and empty- or immuno-NPs for 48 hrs. Numbers indicate band density normalized to β-actin loading control. (**G**) RT-qPCR analysis of IFN pathway and SASP gene expression in *KPC1* PDAC cells treated as in (F) (*n* = 3 samples per group). A.U., arbitrary units. (**H**) Representative histograms (left) and quantification of MHC-I (H-2k^b^) MFI (right) on *KPC1* PDAC cells treated as in (F) (*n* = 6 samples per group). (**I**) RT-qPCR analysis of antigen presentation/processing gene expression in *KPC1* transplant PDAC tumors from mice treated as in (B) (*n* = 3 mice per group). A.U., arbitrary units. Error bars, mean ± SEM. *P* values were calculated using two-tailed, unpaired Student’s t-test. **** P <0.0001, *** P <0.001, ** P <0.01, * P <0.05. n.s., not significant.

Given the unexpected induction of IFNβ within PDAC tumor cells, we investigated whether T/P and immuno-NP treatment also synergized in a tumor cell autonomous manner to further enhance pro-inflammatory SASP signaling. Combined treatment of murine *KPC* PDAC tumor cells in culture led to enhanced phosphorylation of STING and downstream TBK1 and IRF3, indicating activation of this pathway, as well as p65 that we have shown to be a master transcriptional regulator of the pro-inflammatory SASP (*12, 33*) (Fig. 2, E and F). Whereas immuno-NPs had little effect on their own, their combination with T/P produced significant induction of interferon genes and SASP cytokines and chemokines in both murine *KPC* as well as human PANC-1 PDAC tumor cells (Fig. 2G and fig. S2B). Moreover, combined T/P and immuno-NP treatment also significantly increased MHC-I expression on tumor cells (Fig. 2H). An increase in antigen presentation/processing gene expression was also observed in bulk PDAC tumors treated with the combination *in vivo* (Fig. 2I). Taken together, these findings demonstrate that senescence-inducing T/P and immuno-NP treatment synergize through both tumor cell autonomous and non-cell autonomous molecular mechanisms to enhance interferon and pro-inflammatory cytokine production and antigen presentation in the PDAC TME.

### Immuno-NP and T/P treatment activates cytotoxic NK and T cell immunity in PDAC models

Given their synergistic effects on antigen presentation and interferon and cytokine signaling in the PDAC TME, we next investigated the impact of immuno-NP and T/P therapy on innate and adaptive immune responses in *KPC* PDAC transplant models and GEMMs. Similar to our previous findings, while a 2-week T/P treatment increased CD4^+^ and CD8^+^ T cell numbers and a single dose of immuno-NPs enhanced NK cell accumulation and proliferation (as marked by CD69), neither of these single treatment arms alone was able to induce robust NK and T cell cytotoxicity as assessed by expression of the degranulation marker Granzyme B (GZMB) (Fig. 3, A to E). In contrast, combined immuno-NP and T/P treatment led not only to a further increase in CD4^+^ and CD8^+^ T cell and NK cell numbers and infiltration within tumor areas compared to each single treatment regimen, but also enhanced expression of both CD69 and GZMB activation markers on CD8^+^ T cells (Fig. 3, A to E). Moreover, though total numbers of CD4^+^ T cells increased, FOXP3^+^ regulatory T cells (Tregs) that act to inhibit cytotoxic CD8^+^ T cell activity were severely reduced following combined treatment (Fig. S3A). In addition, the amount of mature and activated TNFα^+^ macrophages and DCs, as well as MHC-II^+^ DCs that present antigen to T cells, also expanded following dual NP and T/P therapy (Fig. 3, F to H, and fig. S3B). Thus, combined immuno-NP and T/P treatment leads to reduced Treg numbers, increased antigen-presenting DCs, and significantly enhanced NK and CD8^+^ T cell accumulation and activation in the PDAC TME.

**Fig. 3.**
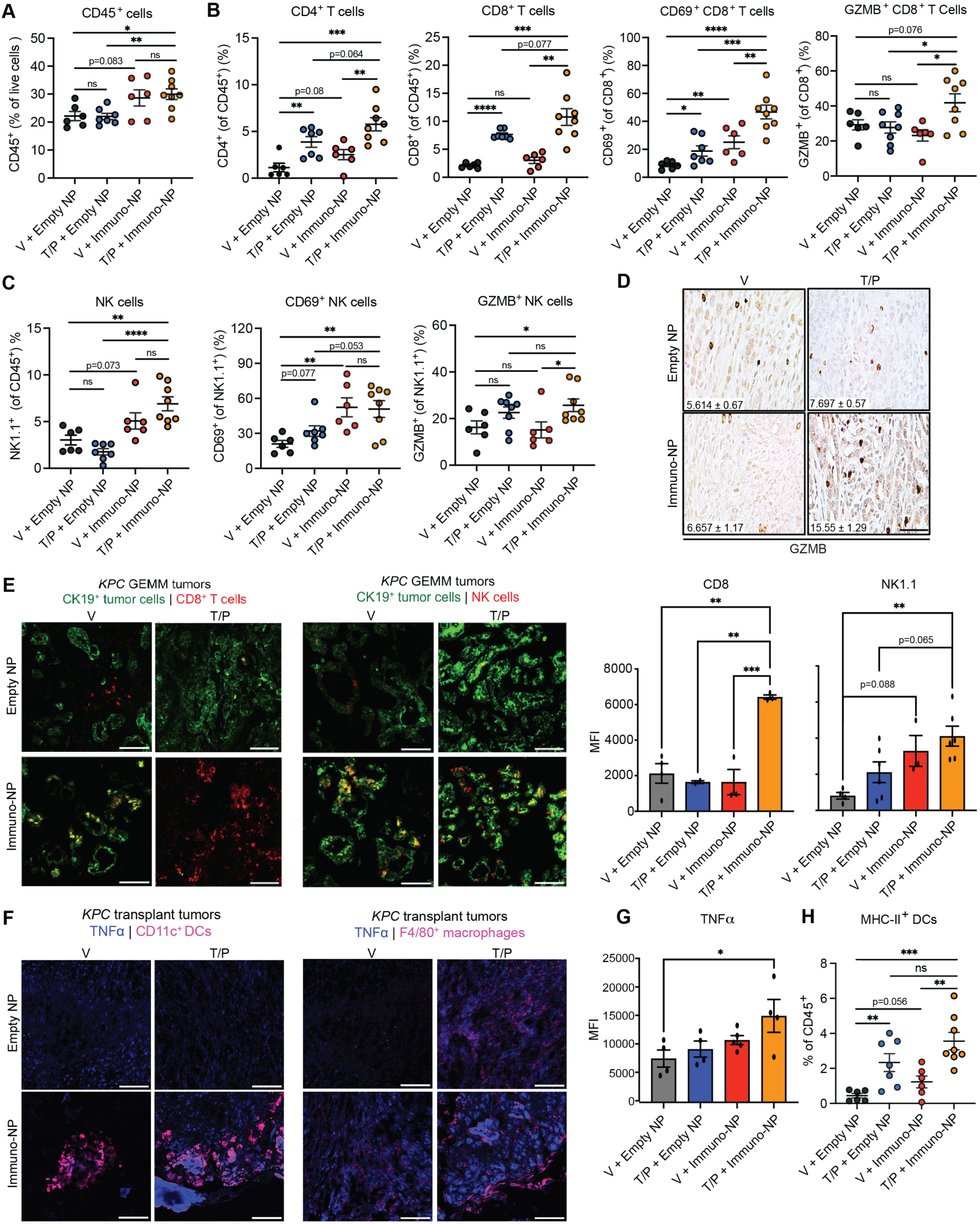
Combinatorial immuno-NP and T/P treatment activates NK and CD8^+^ T cell immunity in PDAC. (**A** to **C**) Flow cytometry analysis of total CD45^+^ immune cells (**A**), T cell numbers and activation markers (**B**), and NK cell numbers and activation markers (**C**) in *KPC1* orthotopic transplant PDAC tumors from mice treated with vehicle or trametinib (1 mg/kg) and palbociclib (100 mg/kg) for 2 weeks and empty- or immuno-NPs for 48 hrs (*n* = 6 to 8 mice per group). (**D**) Immunohistochemical (IHC) staining of *KPC1* orthotopic transplant PDAC tumors from mice treated as in (A). Quantification of the number of degranulating Granzyme B (GZMB)^+^ cells per field is shown inset (*n* = 3 to 6 mice per group). Scale bar, 50μm. (**E**) IF staining of PDAC tumors from *KPC* GEMM mice treated as in (A) (left). Quantification of NK1.1^+^ NK cell and CD8^+^ T cell MFI is shown on right (*n* = 3 mice per group). Scale bars, 100 μm. (**F**) IF staining for TNFα expression in CD11c^+^ DCs (left) and F4/80^+^ macrophages (right) in PDAC tumors from *KPC1* transplant mice treated as in (A). Scale bars, 100 μm. (**G**) Quantification of combined TNFα MFI in macrophages and DCs from IF staining in (F) (*n* = 3 mice per group). (**H**) Flow cytometry analysis of MHC-II^+^ DCs in *KPC1* transplant PDAC tumors from mice treated as in (A) (*n* = 6 to 8 mice per group). Error bars, mean ± SEM. *P* values were calculated using two-tailed, unpaired Student’s t-test. **** P <0.0001, *** P <0.001, ** P <0.01, * P <0.05. n.s., not significant.

### Immuno-NP and T/P therapy leads to tumor regressions and long-term survival in preclinical PDAC models

Combined immuno-NP and T/P treatment also mediated profound short- and long-term anti-tumor responses. Two-week treatment of PDAC-bearing *KPC* transplant mice with daily T/P and weekly immuno-NP administration significantly reduced tumor growth compared to immuno-NP treatment alone (Fig. 4A). Moreover, T/P treatment for 2 weeks followed by a single dose of immuno-NPs produced large areas of tumor necrosis within just 48 hrs (Fig. 4B). This increased tumor control following dual immuno-NP and T/P therapy led to a significant improvement in the overall survival of PDAC-bearing *KPC* transplant mice compared to either single therapy alone following continuous treatment (Fig. 4C). The anti-tumor responses were even more striking in autochthonous *KPC* GEMM mice, where combined immuno-NP and T/P treatment led to tumor necrosis and shrinkage in 8/9 as compared to 3/7 or 0/5 animals treated with immuno-NP or T/P alone, respectively (Fig. 4, D and E). These tumor regressions not only led to significant increases in long-term survival, but, remarkably, 20% of mice treated with immuno-NPs and T/P had complete tumor responses that were maintained even after treatment was stopped (Fig. 4F). Together, these results demonstrate that combined immuno-NP and T/P therapy can produce long-term tumor control and even curative responses in preclinical PDAC models.

**Fig. 4.**
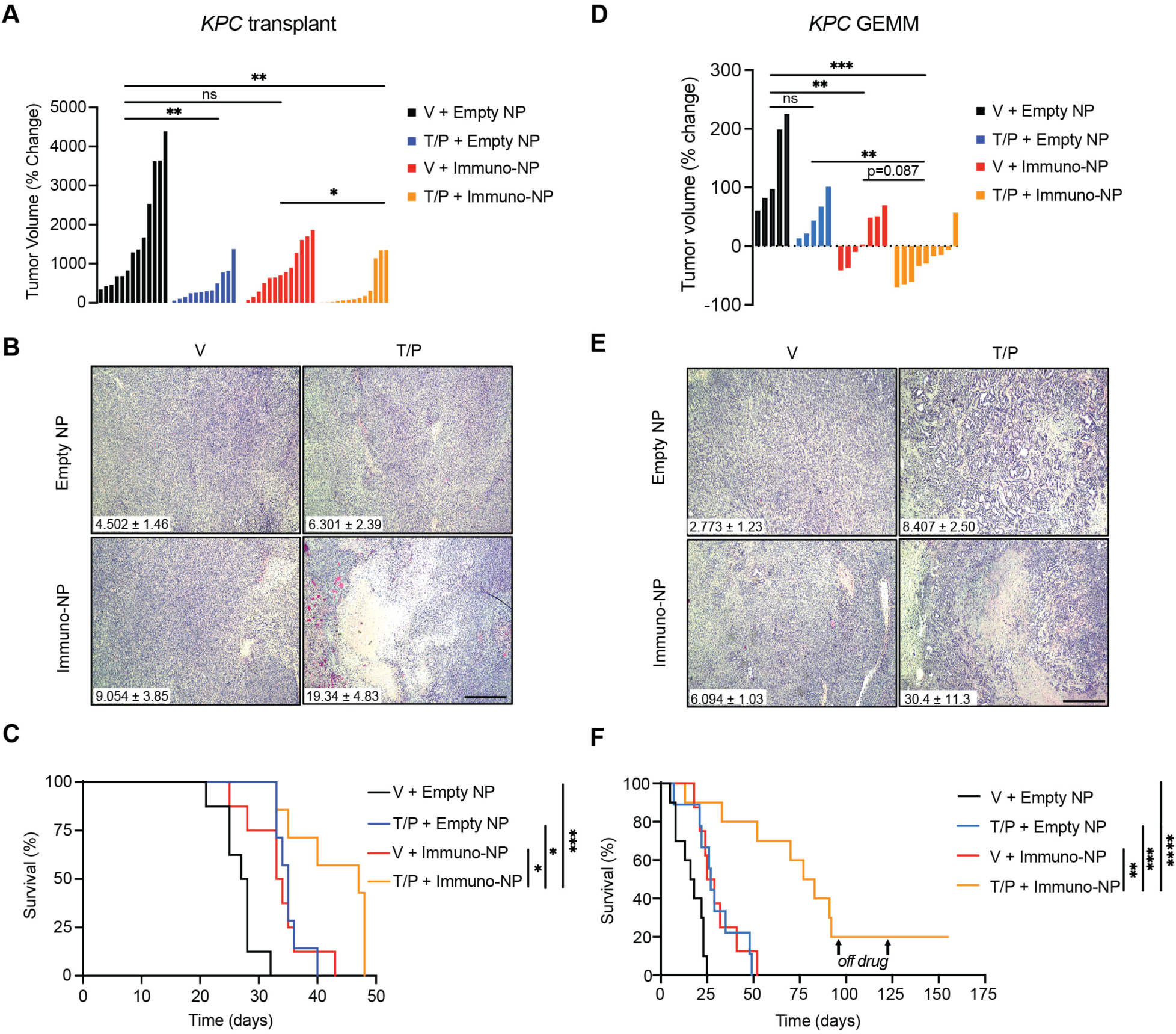
Immuno-NP and T/P regimens produce tumor control and substantially increase overall survival in preclinical PDAC models. (**A**) Waterfall plot of the response of *KPC1* transplant PDAC tumors after treatment with vehicle or trametinib (1 mg/kg) and palbociclib (100 mg/kg) 4 times per week and empty- or immuno-NPs weekly for 2 weeks (*n* = 12 to 13 mice per group). (**B**) H&E staining of *KPC1* transplant PDAC tumors from mice treated with vehicle or trametinib (1 mg/kg) and palbociclib (100 mg/kg) for 2 weeks and empty- or immuno-NPs for 48 hrs. Quantification of percent of tumor area covered in necrosis is shown inset (*n* = 5 to 6 mice per group). Scale bar, 500μm. (**C**) Kaplan-Meier survival curve of mice harboring *KPC1* transplant PDAC tumors treated with vehicle or trametinib (1 mg/kg) and palbociclib (100 mg/kg) 4 times per week and empty- or immuno-NPs weekly (*n* = 7 to 8 mice per group). (**D**) Waterfall plot of the response of *KPC* GEMM PDAC tumors to treatment as in (A) (*n* = 5 to 9 mice per group). (**E**) H&E staining of *KPC* GEMM PDAC tumors from mice treated as in (B). Quantification of percent of tumor area covered in necrosis is shown inset (*n* = 4 to 7 mice per group). Scale bar, 500μm. (**F**) Kaplan-Meier survival curve of PDAC-bearing *KPC* GEMM animals treated as in (C) (*n* = 8 to 10 mice per group). Arrows indicate when mice were taken off of treatment. Error bars, mean ± SEM. *P* values were calculated using two-tailed, unpaired Student’s t-test (A and D) or log-rank test (C and F). **** P <0.0001, *** P <0.001, ** P <0.01, * P <0.05. n.s., not significant.

### Type I IFN-mediated NK and CD8^+^ T cell immunity drives treatment efficacy

To assess whether the tumor responses and survival benefit observed upon dual immuno-NP and T/P treatment were dependent on cytotoxic lymphocyte immunity, we used monoclonal antibodies targeting NK1.1 (PK136) and CD8 (2.43) to deplete NK and cytotoxic T cells, respectively. CD8^+^ T cell, and to a lesser but still significant extent NK cell depletion, both resulted in increased tumor growth and reduced the survival of PDAC-bearing *KPC* transplant mice treated with combination therapy (Fig. 5, A and B). This suggests that NK and CD8^+^ T cells that we have shown to increase in number and become activated following therapy (Fig. 3, A to E) are necessary for its anti-tumor efficacy.

**Fig. 5.**
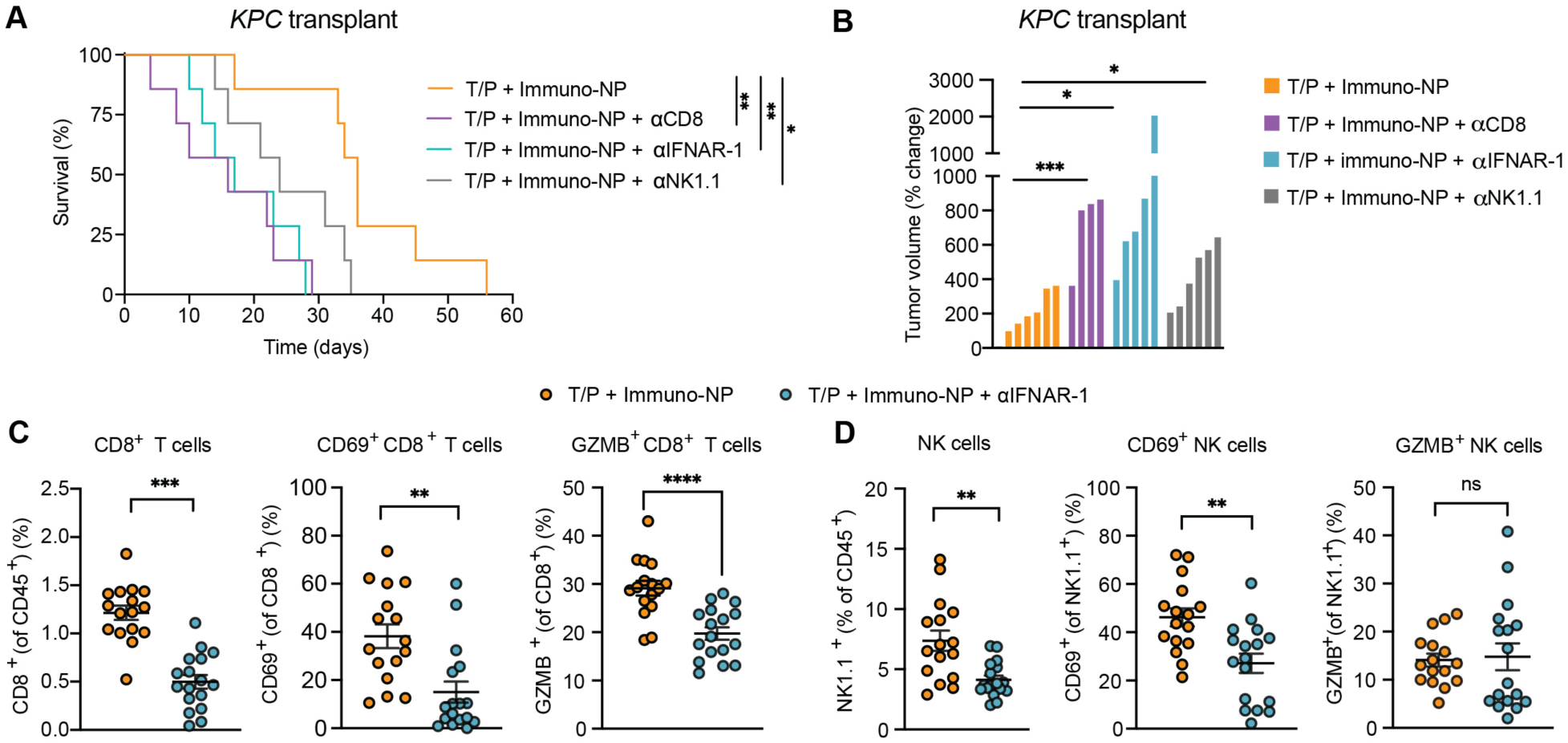
Immuno-NP and T/P therapy efficacy driven by IFNAR-dependent NK and CD8^+^ T cell immune surveillance. (**A**) Kaplan-Meier survival curve of mice harboring *KPC1* transplant PDAC tumors treated with trametinib (1 mg/kg) and palbociclib (100 mg/kg) 4 times per week, immuno-NPs weekly, and blocking antibodies against NK1.1 (PK136; 250 μg), CD8 (2.43; 200 μg), or IFNAR-1 (MAR15A3; 200 μg) twice per week (*n* = 7 mice per group). (**B**) Waterfall plot of the response of *KPC1* transplant PDAC tumors to 2 weeks of treatment as in (A) (*n* = 4 to 7 mice per group). (**C** to **D**) Flow cytometry analysis of CD8^+^ T cell (**C**) and NK cell (**D**) numbers and activation markers in *KPC1* transplant PDAC tumors from mice treated with trametinib (1 mg/kg) and palbociclib (100 mg/kg) 4 times per week, immuno-NPs weekly, and neutralizing antibodies against IFNAR-1 (MAR15A3; 200 μg) administered twice per week for 2 weeks (*n* = 16 to 17 mice per group). Error bars, mean ± SEM. *P* values were calculated using log-rank test (A) or two-tailed, unpaired Student’s t-test (B to D). **** P <0.0001, *** P <0.001, ** P <0.01, * P <0.05. n.s., not significant.

IFNβ that is synergistically induced in tumor and immune cells following immuno-NP and T/P treatment (Fig. 2, C and D) can activate cytotoxic NK and T cell immunity by binding to the Type I interferon receptor, IFNAR, expressed on these cells (*34*). To explore the role of Type I interferon signaling in anti-tumor NK and T cell immunity following therapy, we also treated mice with an IFNAR-1 neutralizing antibody. IFNAR blockade not only significantly reversed NK and CD8^+^ T cell infiltration and activation induced by immuno-NP and T/P treatment, but also reduced the survival benefit of treatment to a similar extent as CD8 depletion alone (Fig. 5, A to C). Collectively, these findings demonstrate that Type I interferon signaling induced upon combined immuno-NP and T/P treatment leads to anti-tumor NK and CD8^+^ T cell responses that mediate immunological tumor control in PDAC models.

### STING and TLR4-driven Type I IFN signaling associated with NK and T cell signatures in human PDAC

To determine whether STING and TLR4 activity and downstream IFN signaling are associated with NK and T cell immunity in human PDAC, we performed analysis on patient samples from two separate published transcriptomic datasets (*35, 36*). STING (*TMEM173*) and *TLR4* expression in patient tumors positively correlated with expression of NK and T cell signature genes in both datasets (Fig. 6). Moreover, high expression of downstream STING and TLR4 pathway components, as well as IRF3 target and Type I IFN signaling genes were also associated with increased NK and T cell transcripts. Taken together with our functional experiments in preclinical PDAC mouse models, these results suggest that potent activation of downstream Type I IFN signaling could be a promising therapeutic avenue to reactivate anti-tumor immunity in PDAC patients.

**Figure 6.**
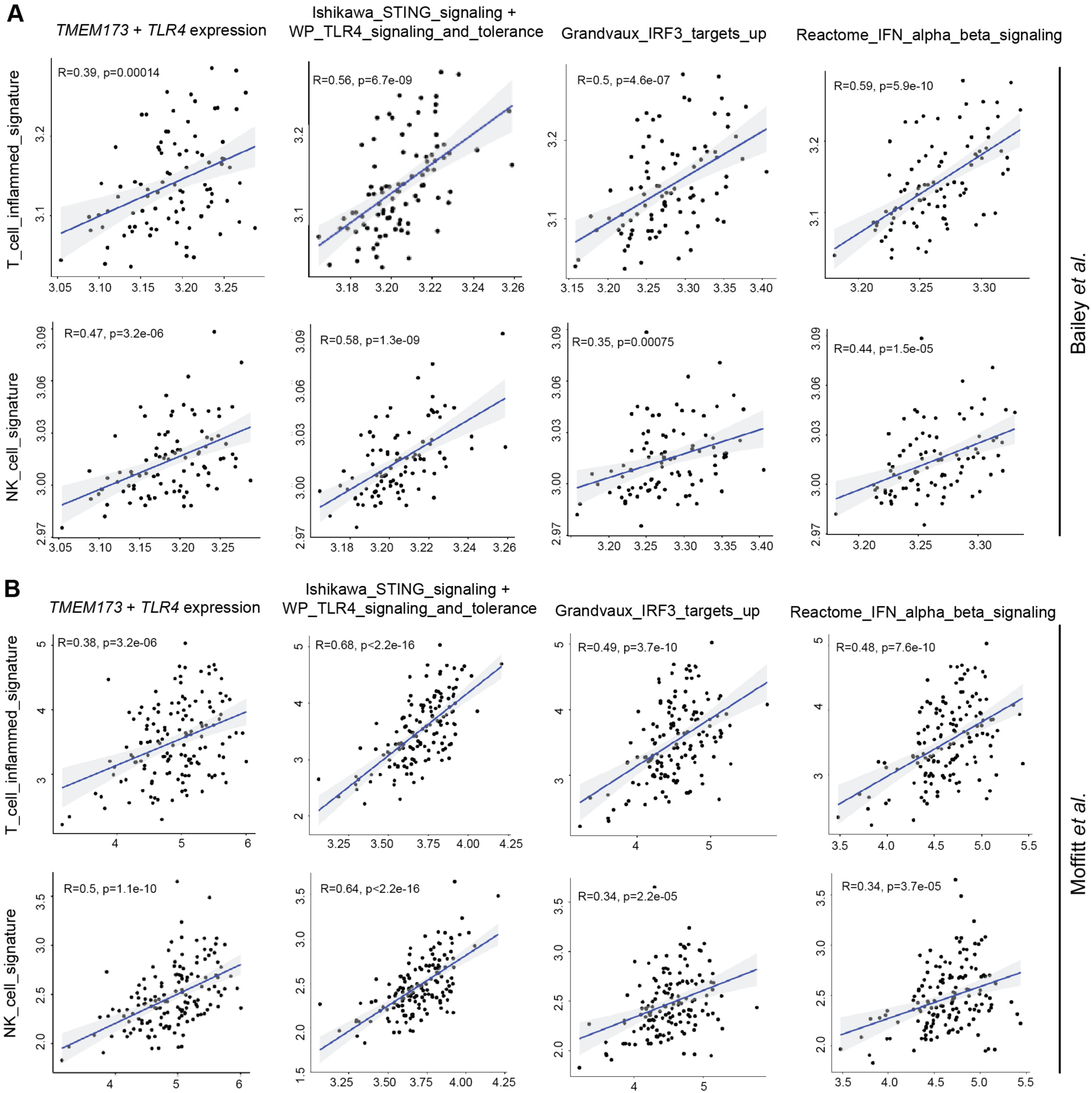
STING and TLR4 expression and Type I interferon signaling correlate with NK and T cell immunity in human PDAC. (**A** to **B**) Pearson’s correlation analysis plots comparing NK and T cell signatures with expression of STING (*TMEM174*), *TLR4*, and downstream interferon signaling pathway genes in human PDAC transcriptomic data from Bailey *et al.* (*35*) (**A**) and Moffitt *et al.* (*36*) (**B**) (*n* = 91 to 145 samples). Pearson’s correlation coefficient (R) values are displayed. *P* values were calculated using a two-tailed, unpaired Student’s *t*-test.

## DISCUSSION

The pancreatic TME presents multiple immune suppressive hurdles that must be overcome for effective immunotherapy, including (a) a desmoplastic stroma contributing to physical exclusion and chemical inhibition of immune cells, (b) a poorly vascularized matrix leading to poor delivery of drugs and infiltration of peripheral immune cells, (c) a lack of cytotoxic NK and T lymphocytes that drive tumor eradication, (d) suppressive myeloid populations that inhibit lymphocyte activation, (e) few DCs to present antigen to T cells, and (f) low neo-antigen loads and dysfunctional antigen presentation circuitry in tumor cells allowing them to escape immune detection (*2, 5*). Though other groups have started to show promising early clinical results with neo-antigen vaccines (*37*), myeloid reprogramming (*38, 39*), and stromal remodeling agents (*40, 41*) that target some of these immune suppressive mechanisms, none of these therapies have yet to be clinically approved. Here we designed a novel multi-pronged approach combining localized innate immune agonist and systemic tumor-targeting senescence-inducing therapies to target many of the immune suppressive mechanisms in PDAC simultaneously. As we had previously shown that therapy-induced senescence with T/P treatment produces increased vascularization, tumor antigen presentation, and CD8^+^ cell infiltration in the PDAC TME (*13*), and immuno-NPs loaded with STING and TLR4 agonists can be taken up by APCs that promote local NK and T cell activation (*27, 30–32*), we hypothesized these two therapy modalities would effectively combine to sustain cytotoxic T cell immunity against PDAC. Indeed, we found remarkable synergy between T/P and immuno-NP therapies, which together led to increased immune agonist uptake and activity, Type I interferon and cytokine signaling, antigen presentation by both tumor cells and APCs, and potent and sustained CD8^+^ T cell activation that culminated in durable and even curative PDAC tumor responses in preclinical animal models. Mechanistically, Type I interferon signaling was key to these therapeutic responses by increasing tumor immunogenicity and coordinating an orchestrated innate and adaptive immune attack, which may be pivotal to mediate immune control of PDAC.

STING agonists have been actively pursued as immune oncology agents in PDAC as well as other solid tumor malignancies as a means to activate Type I interferon signaling that is critical for antigen presentation and productive anti-tumor innate and adaptive immune responses (*16, 21–23*). However, to date STING agonists have yet to show effective clinical utility in cancer in part through their limited cellular uptake, inflammatory toxicities associated with systemic administration, and long-term effects on T cell viability and exhaustion (*24–26*). A major innovation of our study is the ability to systemically deliver STING agonists locally to diverse cell types within the hard-to-penetrate PDAC TME through the design of lipid nanoparticles (NPs) engineered to preferentially deposit in the “leaky” tumor endothelium. Moreover, our nanomaterials-based drug delivery approach enables us to effectively and safely co-deliver physically and chemically distinct STING (cdGMP) and TLR4 (MPLA) agonists, which we and others have shown can together drive robust downstream IRF3 and subsequent Type I interferon signaling (*27, 30, 31, 42*) without systemic toxicities. Remarkably, T/P pre-treatment, likely through the vascular remodeling capabilities of its pro-angiogenic SASP, further increased the deposition of NPs in the PDAC TME. As such, our engineering approach augmented through senescence-induced vascular remodeling can overcome some of the drug delivery challenges that have been a major limitation to the effective treatment of PDAC and clinical development of STING agonists.

Unexpectedly, we found that immuno-NPs were not only taken up by APCs in the perivascular regions of the PDAC TME, but also by tumor cells themselves, where they synergized with T/P-induced senescence to enhance the pro-inflammatory SASP in a tumor cell autonomous manner. The SASP can be a double-edged sword in cancer, with some SASPs promoting anti-tumor immune surveillance, while others pro-tumor immune suppression, and its context-dependent regulation is only beginning to be understood (*14, 43*). Indeed, we have previously shown that whereas T/P-induced senescence drives immune-mediated tumor regressions in KRAS mutant lung cancers, it does not produce the same immune responses or tumor control KRAS mutant PDAC (*12, 13, 15*). Our work here demonstrates a new means of SASP regulation by which STING/TLR4-mediated signaling enhances not only IFNβ production but also a slew of pro-inflammatory SASP cytokines and chemokines that are normally repressed in the PDAC tumors even after T/P treatment to achieve NK and CD8^+^ T cell immune control. Though we demonstrate that Type I interferon signaling through its receptor IFNAR is critical for cytotoxic lymphocyte immunity with this combination therapy, it is possible that other SASP-associated chemokines and cytokines that are also synergistically enhanced and that we have previously implicated in activating NK and T cell immunity, including CCL2, CXCL9/10, and IL-12/-15/-18, could also contribute to immune responses to therapy. In addition, enhanced MHC-I expression on tumor cells and MHC-II expression on DCs, presumably induced through IFNAR signaling, could also contribute to enhanced CD8^+^ T cell responses to treatment, and if so may suggest rationale combinations with neo-antigen or DC vaccines to sustain durable T cell responses against PDAC.

Our study has several limitations. Given the diverse cell types and mechanisms of immune suppression targeted by these therapies, moving forward single cell sequencing modalities will be critical to understanding the key tumor, immune, and stromal (e.g. endothelial cells, fibroblasts) cell types and mechanisms that T/P and immuno-NP treatment act on to elicit anti-tumor immunity. Though NPs effectively deliver cargo and induce IFNβ in both tumor cells and APCs, it is unclear whether targeting one or both cell types is necessary to achieve anti-tumor immunity. Moreover, many different immune cell subsets also express IFNAR and can respond to Type I interferons beyond just NK and CD8^+^ T cells (*34*), including suppressive myeloid cells and regulatory T cells (Tregs) that were diminished following treatment and whose targeting could indirectly enhance cytotoxic lymphocyte activity. In the future, transgenic models with cell type-restricted IRF3 or IFNAR knockout could be employed to tease apart the IFN signaling crosstalk that ultimately drives cytotoxic NK and CD8^+^ T cell immunity in our system (*44, 45*). As our experiments focused primarily on short-term treatment effects, further studies are needed to assess the long-term effects of IFN signaling, which could lead to eventual T cell exhaustion (*46–49*), on PDAC immune responses, and to best optimize dose and scheduling of these regimens. Provided the tunability of our nanomaterials engineering approaches, we can also add targeting peptides to NPs to target them to specific cell types in the PDAC TME, and design them to encapsulate T/P to further optimize their tumor delivery and on-target effects.

Immuno-NP and T/P treatment in human PDAC cells and analyses of patient PDAC samples suggests that these therapies can activate potent Type I interferon signaling, and that this is associated with enhanced NK and T cell immunity in human PDAC, highlighting the translational potential of our approach. Given differences in STING protein structure between mice and humans (*18*), our modular lipid-based NP approach can be adapted to substitute other STING agonists (e.g. cGAMP, diABZI) that are may be more suitable for human testing. As our previous work demonstrated that FDA-approved trametinib, palbociclib, and anti-PD-1 antibodies can synergize effectively in PDAC models (*13*), future studies and trials could involve the combination of immuno-NPs and T/P with anti-PD-1 ICB, which has yet to be effective on its own in PDAC, to determine if these regimens could have clinical utility. Collectively, our results broadly suggest that engineering approaches to target multiple cell types and immune suppressive barriers through induction of Type I interferon signaling in the PDAC TME could pave the way for coordinated innate and adaptive immune responses to achieve immunotherapy success that has thus far been elusive for PDAC patients.

## MATERIALS AND METHODS

### Study design

Sample sizes were determined based on those reported in previous publications (*12, 13, 15*) and no statistical method was used to predetermine sample size. The indicated sample size (*n*) represents biological replicates. All experiments were repeated independently 2-3 times. All samples that met proper experimental conditions were included in the analysis. For *in vivo* experiments, mice were randomized based on tumor burden as assessed by ultrasound imaging to achieve equal tumor volume between experimental groups. For *in vitro* experiments sample allocation was performed randomly. Data collection and analysis were not performed in a blinded manner.

### Nanoparticle synthesis and characterization

Dual agonist immuno-NPs were synthesized by pulsed ultrasonication. Equimolar amounts of DOPC (33.5 mol% 1,2-dioleoyl-sn-glycero-3-phophocholine, Avanti) and DSPC (33.5 mol% 1,2-distearoyl-sn-glycero-3-phosphocholine, Avanti) as well as DOPG (20 mol% 1,2-dioleoyl-sn-glycero-3-phospho-(1’-rac-glycerol), Avanti) were prepared in chloroform, along with 10 mol% cholesterol and 3 mol% mPEG2000-DSPE [methoxy-poly(ethyleneglycol)-2000 1,2-distearoyl-sn-glycero-3-phophoethanolamine-N, Laysan Bio]. MPLA (Sigma-Aldrich) was added to lipids in chloroform and dried to form lipid films. For some experiments a lipophilic fluorescent Di tracer (i.e. DiI, DiD) was also added to the bilayer at 0.1 mol%. Films were then rehydrated in PBS containing cdGMP (Invivogen), heated to 60°C for 1h, with vortexing every 10 minutes for 30s intervals. Samples were ultrasonicated on ice using 30s cycles with pulsing at 20% amplitude for 20s followed by a 10s pause for a total of 5 minutes. Immuno-NPs were then dialyzed for 1h against sterile PBS and stored immediately at 4°C. Dynamic light scattering (DLS) and zeta potential were used to measure immuno-NP hydrodynamic size and surface, respectively, using a Malvern Zetasizer. A commercially available cdGMP detection kit (Lucerna Technologies) was used to quantify cdGMP encapsulation. Empty NPs lacking MPLA and cdGMP were used as a vehicle control.

### Immuno-NP safety studies

Wild-type (WT) C57BL/6 mice were treated weekly with immuno-NPs containing 7 µg of cdGMP and MPLA each by intravenous (i.v.) injection for 3 consecutive weeks. Animal weight was recorded weekly. Blood was collected in heparinized tubes and plasma separated *via* centrifugation at 1,500xg for 20 min at 4°C following 3-week treatment. Plasma samples were sent to the UMass Chan Medical School Analytical Core for analysis of ALT/AST levels. Liver toxicity was assessed histopathologically using H&E-stained sections.

### Animal studies

All mouse experiments in this study were approved by the University of Massachusetts Chan Medical School Internal Animal Care and Use Committee. Mice were maintained under specific pathogen-free conditions, and food and water were provided ad libitum. C57BL/6 mice for transplantation models were purchased from Charles River Laboratories and *KPC* GEMM mice were bred in-house.

### Pancreas orthotopic transplant models

5×10^4^ *KPC1* cells were resuspended in 25μl of Matrigel (Matrigel, BD) diluted 1:1 with cold advanced DMEM/F12 media and transplanted into the pancreas of 8–10-week-old C57BL/6 female mice. After administering anesthesia using 2-3% isoflurane, an incision was performed on the left side of the abdomen. Subsequently, the cell suspension was injected into the tail region of the pancreas using a Hamilton Syringe. The injection’s success was confirmed by the presence of a fluid bubble without any indications of leakage into the abdominal cavity. The abdominal wall was closed using an absorbable Vicryl suture (Ethicon), and the skin was secured with wound clips (CellPoint Scientific Inc.). Mice were then monitored for tumor development using ultrasound imaging. One week after transplantation, mice were randomized into different treatment groups based on tumor volume. Following sacrifice, a portion of the pancreas tumor tissue was preserved in 10% formalin for fixation, while others were used for OCT frozen blocks and flow cytometry analysis.

### *KPC* genetically engineered mouse model (GEMM)

*P48-Cre; Kras^LSL-G12D/wt^; Trp53^fl/wt^* (*KPC*) GEMMs were generated by interbreeding *P48-Cre, Kras^LSL-G12D/wt^*, and *Trp53^fl/fl^* strains on a C57BL/6 background. Tumor development was monitored using ultrasound imaging. Once tumors reached approximately 50 mm^3^ in volume, the mice were enrolled and randomized into different treatment groups based on tumor volume. After sacrificing the mice, pancreatic tumor tissue was divided for 10% formalin fixation for immunohistochemistry (IHC) and OCT frozen blocks for immunofluorescence (IF) assays.

### Drug treatments and neutralizing antibodies

Trametinib was dissolved in a solution containing 0.5% hydroxypropyl methylcellulose and 0.2% Tween-80 and palbociclib in 50 mM sodium lactate buffer (pH 4). PDAC-bearing mice were treated with vehicles or trametinib (1 mg/kg) and palbociclib (100 mg/kg) (LC Laboratories) orally for four consecutive days followed by three days without treatment for two weeks or until survival endpoint. For short-term 48 hr or 2-week treatment studies, PDAC-bearing mice received a single dose of control empty NPs or immuno-NPs carrying 7 µg of cdGMP and MPLA each by intravenous (i.v.) injection, and animals euthanized and tumors harvested 48 hrs after treatment. For toxicity and long-term survival experiments, PDAC-bearing mice received empty or immuno-NPs weekly. To deplete NK or CD8^+^ T cells, mice received intraperitoneal (i.p.) injections of an αNK1.1 (250 μg; PK136, BioXcell) or αCD8 (200 μg; 2.43, BioXcell) antibody twice per week, respectively. To neutralize IFNAR signaling, mice were i.p. injected with an αIFNAR-1 antibody (200 μg; MAR15A3, BioXcell) twice per week. No toxicities (as assessed by weight loss and liver damage) were observed in animals treated with these compounds alone or in combination. Ultrasound imaging was performed every two weeks during the treatment period to monitor changes in PDAC tumor burden.

### Ultrasound imaging

To stage and quantify tumor burden, high-contrast ultrasound imaging was performed using a Vevo 3100 System with a MS250 13- to 24-MHz scanhead (VisualSonics). Tumor volume was analyzed using Vevo LAB software.

### Immunofluorescence (IF)

Fresh tissues were embedded in OCT, frozen, and cut into 5 μm sections (taken from center of the tissue). Tissue sections were placed in humidity chambers for staining. In brief, samples were washed 3x with PBS prior to fixation with 2% PFA for 1h at RT. PFA was removed and protein blocking solution (4% goat serum, 0.5% triton-X in PBS) was added for 20min at RT. The following primary antibodies diluted in protein blocking solution were added to the tissue and incubated overnight at 4°C: CK19 (1:200; TROMA-III, Univ. of Iowa Developmental Studies Hybridoma Bank), CD31 (1:100; polyclonal, Thermo Fisher), IFNβ (1:100; polyclonal, Thermo Fisher), CD11c (1:100; N418, Thermo Fisher), F4/80 (1:100; A3-1, Thermo Fisher), TNFα (1:100; polyclonal, Thermo Fisher), CD8a (1:100; 53-6.7, Thermo Fisher), and NK1.1 (1:100; polyclonal, Thermo Fisher). Tissues were then washed 3x with PBS and secondary Alexa Fluor 405, 488, 568, or 647 dye-conjugated antibodies (Thermo Fisher) were added diluted 1:150 in protein blocking solution for 1h at RT. Tissue sections were washed 3x with PBS and mounting media with or without DAPI (Vectashield) was added prior to applying a glass coverslip. Images were obtained using a Nikon A1 confocal microscope or Leica Thunder Live Cell and 3D Assay Imager. Fluorescence was analyzed and quantified using Fiji/ImageJ.

### Immunohistochemistry (IHC)

Tissues were fixed overnight in 10% formalin, embedded in paraffin, and cut into 5 μm sections. Hematoxylin and eosin (H&E) and immunohistochemical staining were performed using standard protocols. For immunohistochemistry, sections were deparaffinized, rehydrated with decreasing concentrations of ethanol in water, and boiled in a pressure cooker for 20 minutes in 10 mM citrate buffer (pH 6.0). Endogenous peroxidases were quenched by incubating the slides in 3% hydrogen peroxide for 20 min. The sections were then washed 2x with PBS and the following primary antibodies were incubated overnight at 4°C: FOXP3 (1:100; FKJ-16s, eBioscience) and Granzyme B (GZMB) (1:100; AB4059, Abcam). HRP-conjugated secondary antibodies (Vectastain Elite ABC-HRP Kits: Rat, PK-6104; Rabbit, PK-6101) were applied for 30 minutes and visualized with DAB (Vector Laboratories; SK-4100).

For quantification of FOXP3^+^ Tregs and GZMB^+^ immune cells, 5-10 high power 20x fields per section were counted and averaged using ImageJ software. Tumor necrosis was assessed by quantifying the percentage of total PDAC tumor area covered in necrotic tissue from H&E-stained sections using ImageJ software.

### Cell lines and *in vitro* drug treatments

PANC-1 and 293T cells were obtained from the American Type Culture Collection (ATCC). The murine *KPC1* PDAC cell line was generated as described previously (*13*). *KPC1* cells were transduced with an MSCV-luciferase (luc)-IRES-GFP retroviral construct to visualize and track tumor cells following *in vivo* transplantation. Retroviruses were produced by co-transfecting Gag-Pol expressing 293 T cells with the appropriate expression and envelope vectors (VSV-G). After transduction, the cells were purified by FACS sorting the GFP^+^ population using a FACSAria (BD Biosciences). All cells were cultured in a humidified incubator at 37°C with 5% CO_2_ and grown in DMEM supplemented with 10% FBS and 100 IU/ml penicillin/streptomycin (P/S). *KPC*1 cells were grown on culture dishes coated with 100 µg/ml collagen (PureCol) (5005; Advanced Biomatrix). All cell lines used tested negative for mycoplasma. Human cell lines were authenticated by their source repository.

Trametinib (S2673) and palbociclib (S1116) were purchased from Selleck Chemicals and MedChemExpress, dissolved in DMSO (vehicle) to obtain 10mM stock solutions, and stored at - 80°C for *in vitro* studies. Nanoparticles were synthesized using identical methods to those used for *in vivo* studies as described above. Human and mouse PDAC cells were treated with 25 nM trametinib and 500 nM palbociclib for 7 days and received 1 dose of empty or immuno-NPs (carrying 7 µg each of cdGMP and MPLA) on day 5 prior to their harvesting 48 hrs later.

### qRT-PCR

Total RNA was extracted from *KPC* cell lines or bulk PDAC tumor tissue using the RNeasy Mini Kit (Qiagen). Complementary DNA (cDNA) was obtained using TaqMan reverse transcription reagents (Applied Biosystems). Real time qPCR was performed in triplicate using Power SYBR™ Green PCR Master Mix (Applied Biosystems) on the StepOnePlus RealTime PCR System (Applied Biosciences). The comparative CT method (2^−ΔΔCT^) was used to determine fold differences between the target gene and the reference gene *GAPDH*. Primer sequences are listed in Table S1.

### Immunoblotting

Whole cell protein lysates were extracted using RIPA buffer (Cell signaling) supplemented with phosphatase inhibitors (5mM sodium fluoride, 1 mM sodium orthovanadate, 1 mM sodium pyrophosphate, 1 mM β-glycerophosphate) and protease inhibitors (Protease Inhibitor Cocktail Tablets, Roche). Protein concentration was determined using a Bradford Protein Assay kit (Biorad). Proteins were separated by SDS-PAGE and transferred to polyvinyl difluoride (PVDF) membranes (Millipore) according to standard protocols. Membranes were blotted with antibodies (1:1,000) against phosphorylated (p)-STING^S365^ (D8F4W), p-IRF3^S396^ (4D4G), p-TBK1^S172^ (D52C2s), and p-p65^S536^ (93H1) from Cell Signaling in 5% milk in TBS blocking buffer. After primary antibody incubation, membranes were probed with an ECL anti-rabbit IgG secondary antibody (1:10,000) from GE Healthcare Life Science and imaged using Chemidoc Molecular Imaging System (BioRad). Protein loading was determined using a monoclonal β-actin antibody directly conjugated to horseradish peroxidase (1:20,000) from Sigma-Aldrich (A3854). Quantitation of western blot band intensity was done using Image J software.

### Flow Cytometry

To assess MHC-I expression on *KPC* cells cultured *in vitro*, drug-treated cells were trypsinized, resuspended in PBS supplemented with 2% FBS, and stained with an H-2k^b^ antibody (AF6-88.5.5.3, eBioscience; 1:200) for 30 minutes on ice. Flow cytometry analysis was conducted using a BD LSR II instrument, and FlowJo software (TreeStar) was used for data analysis.

For *in vivo* sample preparation, pancreatic tumor tissues were isolated and allocated for 10% formalin fixation, OCT frozen blocks, and flow cytometry analysis following 48 hr or 2-week treatment. To generate single cell suspensions for flow cytometry analysis, pancreas tumors were minced into small pieces with scissors, placed in 5 ml of collagenase buffer (1x HBSS with calcium and magnesium, 1 mg/ml Collagenase V, 0.1 mg/ml DNase I) in C tubes, and then processed using program 37C_m_TDK1_1 on a gentleMACS Octo dissociator with heaters (Miltenyi Biotec). The dissociated tissue was passed through a 70 μm cell strainer, centrifuged, and resuspended in PBS supplemented with 2% FBS. Samples were then incubated with the following antibodies for 30 minutes on ice: CD45 AF700 (30-F11; 1:320), NK1.1 BV605 (PK136; 1:200), CD3 BV650 (17A2; 1:300), CD8 PE-Cy7 (53-6.7; 1:400), CD4 PE-Cy5 (GK1.5; 1:200), CD69 APC-Cy7 (H1.2F3; 1:200), F4/80 APC (BM8; 1:200), CD11c BV785 (N418; 1:100), MHC-II (I-A/I-E) PE-Dazzle 594 (MS114.15.2; 1:200), Gr-1 (Ly-6G/Ly-6C) Pacific Blue (RB6-8C5, 1:200), B220 (CD45R) PerCP-Cy5.5 (RA3-6B2; 1:400) (Biolegend); and CD11b (M1/70; 1:1,280) (BD Biosciences). DAPI was used to distinguish live/dead cells, and DiI and DiD fluorophores used to mark NPs. Flow cytometry was performed on an BD LSR II, and CD45^+^ immune cell, CD45^-^ GFP^+^ tumor cell, CD45^-^GFP^-^ stromal cell, CD4^+^ and CD8^+^ CD3^+^ T cell, CD3^-^ NK1.1^+^ NK cell, CD3^-^ B220^+^ B cell, CD11b^+^ F4/80^+^ macrophage, CD11b^+^Gr-1^+^ MDSC, CD11b^-^ CD11c^+^ MHC-II^+^ dendritic cell numbers and expression of activation markers (CD69) and NP fluorophores (DiI/DiD) were analyzed using FlowJo (TreeStar).

To analyze Granzyme B (GZMB) expression in NK and T cells, single cell suspensions from tumor tissue were resuspended in RPMI media supplemented with 10% FBS and 100 IU/ml P/S and incubated for 4 hours with PMA (20 ng/ml, Sigma-Aldrich), Ionomycin (1 μg/ml, STEMCELL technologies), and monensin (2 μM, Biolegend) in a humidified incubator at 37°C with 5% CO_2_. Cell surface staining was first performed with CD45 AF700 (30-F11; 1:320), NK1.1 BV605 (PK136; 1:200), CD3 BV650 (17A2; 1:300), CD8 APC-Cy7 (53-6.7; 1:200), and CD4 PE-Cy5 (GK1.5; 1:200) (Biolegend) antibodies. Intracellular staining was then performed using the Foxp3/transcription factor staining buffer set (eBioscience), where cells were fixed, permeabilized, and then stained with a GZMB APC antibody (GB11, Biolegend; 1:100). GZMB expression was evaluated by gating on CD3^-^NK1.1^+^ NK cells and CD3^+^CD8^+^ T cells on an BD LSR II flow cytometer and analyzed using FlowJo (TreeStar) as described above.

### Pearson’s correlation analysis

Gene expression data of primary PDAC patient tumors from two independent studies by Bailey *et al.* (GSE36924)(*35*) and Moffitt *et al.* (GSE71729)(*36*) were downloaded with the GEOquery2 package. Correlation analysis between NK (*50*) and T cell (*51*) gene signatures, STING (*TMEM174*) and *TLR4* gene expression, and STING (*52*), TLR4, IRF3 (*53*), and IFNα/β signaling gene sets was performed using the ggpubr package. Results are presented as Pearson’s correlation coefficient (R) values.

### Statistical analysis

Statistical analyses were performed as described in the corresponding figure legends. Statistical significance was determined by two-sided Student’s *t*-test or log-rank test with Prism 9 software (GraphPad) and R. Values are reported as mean ± standard error of at least 3 independent biological replicates, and sample numbers (*n*) are indicated in the figure legend. Significance was set at *P*<0.05.

## ACKNOWLEDGEMENTS

We thank G. Cottle for technical assistance with i.v. injections. Graphics in Figs. 2A and 2E were created with BioRender.com. The CK19 TROMA-III antibody developed by R. Kemler was obtained from the Developmental Studies Hybridoma Bank, created by the NICHD of the NIH and maintained at The University of Iowa, Department of Biology, Iowa City, IA 52242.

## Funding

UMass Cancer Center Pilot Project Grant (P.U.A., M.R.)

National Cancer Institute (NCI) K22 CA262355 grant (P.U.A.)

National Cancer Institute (NCI) R00 CA241110 grant (M.R.)

National Cancer Institute (NCI) K99 CA252153 grant (J.R.P.)

American Gastroenterological Association (AGA) Bern Schwartz Research Scholar in Pancreatic Cancer (J.R.P.)

## Author Contributions

P.U.A. and M.R. conceived the study, managed the project, designed experiments, interpreted results, and wrote the paper with assistance from all authors. L.C. designed and performed *in vitro* and *in vivo* experiments, analyzed and interpreted results, and wrote the paper. C.F.L., G.I.K., M.L.B., T.E.N., and J.C. synthesized nanoparticles, carried out safety studies, and performed immunofluorescence analysis. K.D.D., C.N.P., and K.C.M. performed and analyzed mouse experiments. J.L. and L.J.Z. performed bioinformatics analysis on human transcriptomic datasets. J.P and J.R.P. performed immunofluorescence analysis. K.A.F. provided reagents and intellectual input on the project.

## Data and materials availability

All reagents and data supporting the findings of this study are available from the corresponding authors upon request.

