## Supplementary Figures and Tables for "Nanoparticle delivery of innate immune agonists combines with senescence-inducing agents to mediate T cell control of pancreatic cancer"

Loretah Chibaya *et al.*

**The PDF file includes:**

Fig. S1 to S3
Table S1

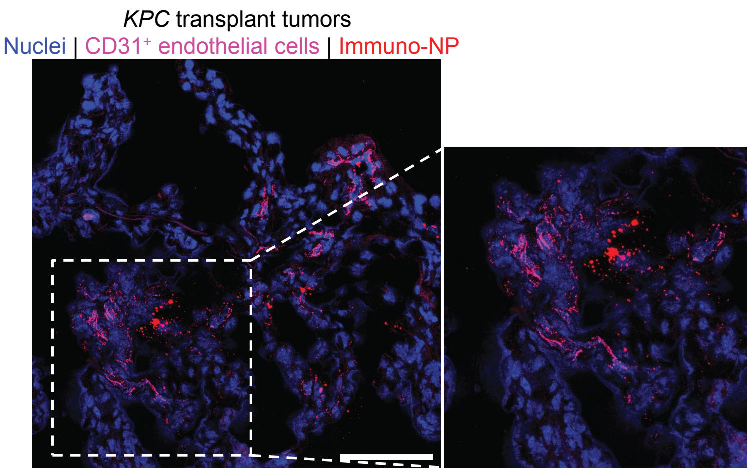

**Fig. S1. Immuno-NPs home to perivascular regions of PDAC TME.** Representative immunofluorescence (IF) staining of *KPC1* orthotopic transplant PDAC tumors for expression of DiI-labeled immuno-NPs in proximity to $CD31^+$ blood vessels. Scale bar, 100 µm.

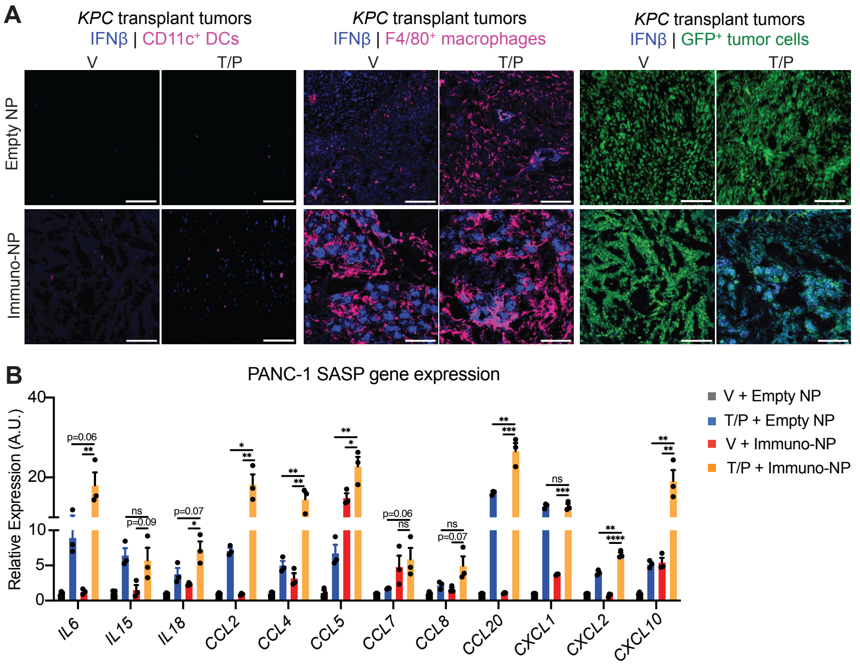

**Fig. S2. T/P and immuno-NP therapy combine to enhance IFNβ expression and SASP production in mouse and human PDAC.** (**A**) Representative IF staining of *KPC1* transplant PDAC tumors from mice treated with vehicle or trametinib (1 mg/kg) and palbociclib (100 mg/kg) for 2 weeks and empty- or immuno-NPs for 48 hrs for expression of IFNβ in DCs ($CD11c^{+}$), macrophages ($F4/80^{+}$), and tumor cells ($GFP^{+}$). Scale bars, 100 μm. (**B**) RT-qPCR analysis of SASP gene expression in PANC-1 human PDAC cells treated with vehicle or trametinib (25 nM) and palbociclib (500 nM) for 1 week and empty- or immuno-NPs for 48 hrs (n = 3 samples per group). A.U., arbitrary units. Error bars, mean ± SEM. *P* values were calculated using two-tailed, unpaired Student's t-test. **** P <0.0001, *** P <0.001, ** P <0.01, * P <0.05. n.s., not significant.

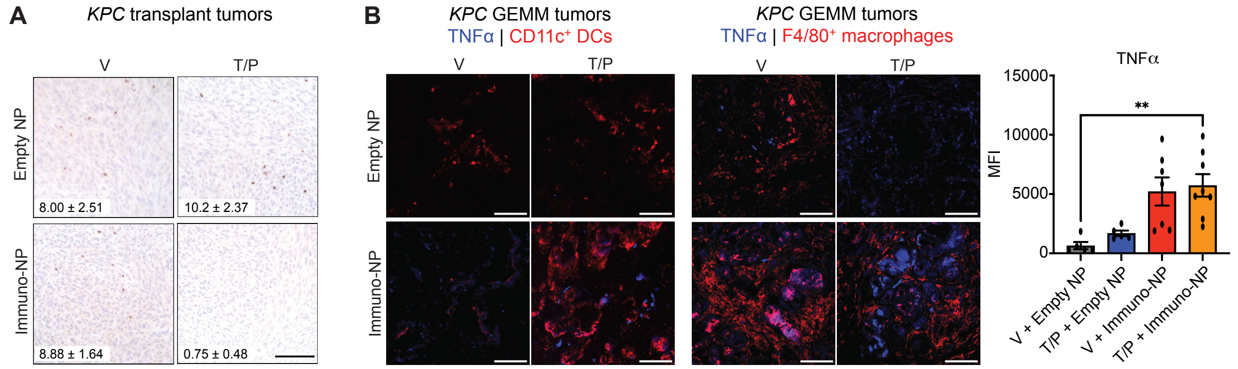

**Fig. S3. Combined T/P and immuno-NP treatment significantly reduces Treg and increases mature macrophage and DC populations in PDAC TME.** (**A**) Immunohistochemical (IHC) staining of *KPC1* orthotopic transplant PDAC tumors from mice treated with vehicle or trametinib (1 mg/kg) and palbociclib (100 mg/kg) for 2 weeks and empty- or immuno-NPs for 48 hrs. Quantification of the number of FOXP3$^+$ Tregs per field is shown inset ($n$ = 4 to 9 mice per group). Scale bar, 100µm. (**B**) IF staining for TNFα expression in CD11c$^+$ DCs (left) and F4/80$^+$ macrophages (right) in PDAC tumors from *KPC GEMM* mice treated as in (A). Quantification of combined TNFα MFI in macrophages and DCs is shown on right ($n$ = 3 mice per group). Scale bar, 100 µm. Error bars, mean ± SEM. *P* values were calculated using two-tailed, unpaired Student's t-test. ** P <0.01.

**Table S1. RT-qPCR primer sequences**

| Mouse | | Human | |
|---|---|---|---|
| Ifnb1 F | CCCTATGGAGATGACGGAGA | IL15 F | AACAGAAGCCAACTGGGTGAATG |
| Ifnb1 R | ACCCAGTGCTGGAGAAATTG | IL15 R | CTCCAAGAGAAAGCACTTCATTGC |
| Tbk1 F | CCGATCTACCCCAGCTCTAA | IL6 F | AGACAGCCACTCACCTCTTCAG |
| Tbk1 R | AAGGCCACCATCCATTGTTA | IL6 R | TTCTGCCAGTGCCTCTTTGCT |
| Irf3 F | CAAGAGGCTTGTGATGGTCA | IL18 F | GATAGCCAGCCTAGAGGTATGG |
| Irf3 R | GCAAGTCCACGGTTTTCAGT | IL18 R | CCTTGATGTTATCAGGAGGATTCA |
| p65 F | AGGCTCCTGTTCGAGTCTCC | CXCL2 F | GGCAGAAAGCTTGTCTCAACCC |
| p65 R | CATAGGTCCTTTTGCGCTTC | CXCL2 R | CTCCTTCAGGAACAGCCACCA |
| Il6 F | TGATGCACTTGCAGAAAACA | CXCL1 F | GCCCAAACCGAAGTCATAGCC |
| Il6 R | ACCAGAGGAAATTTTCAATAGGC | CXCL1 R | ATCCGCCAGCCTCTATCACA |
| Il1b F | TGGACCTTCCAGGATGAGGACA | CXCL10 F | GGTGAGAAGAGATGTCTGAATCC |
| Il1b R | GTTCATCTCGGAGCCTGTAGTG | CXCL10 R | GTCCATCCTTGGAAGCACTGCA |
| Il12 F | ACGAGAGTTGCCTGGCTAG | CCL8 F | TATCCAGAGGCTGGAGAGCTAC |
| Il12 R | CCTCATAGATGCTACCAAGGCAC | CCL8R | TGGAATCCCTGACCCATCTCTC |
| Il15 F | TGCCCTGAACTGCTTTCTCC | CCL20 F | AAGTTGTCTGTGTGCGCAAATCC |
| Il15 R | TCTCCTCCAGCTCCTCACAT | CCL20 R | CCATTCCAGAAAAGCCACAGTTTT |
| Il18 F | GAGGAAATGGATCCACCTGA | CCL4 F | GCTTCCTCGCAACTTTGTGGTAG |
| Il18 R | ATCTTCCTTTTGGCAAGCAA | CCL4 R | GGTCATACACGTACTCCTGGAC |
| Ccl2 F | CCTGCTGTTCACAGTTGCC | CCL7 F | ACAGAAGGACCACCAGTAGCCA |
| Ccl2 R | ATTGGGATCATCTTGCTGGT | CCL7 R | GGTGCTTCATAAAGTCCTGGACC |
| Ccl3 F | CTGCCTGCTGCTTCTCCTAC | CCL2 F | TTCTGTGCCTGCTGCTCATA |
| Ccl3 R | CTTGGACCCAGGTCTCTTTG | CCL2 R | AGCTTCTTTGGGACACTTGC |
| Ccl4 F | GCCCTCTCTCTCCTCTTGCT | CCL5 F | CTGCTGCTTTGCCTACCTCT |
| Ccl4 R | GTCTGCCTCTTTTGGTCAGG | CCL5 R | CGAGTGACAAACACGACTGC |
| Ccl5 F | GTGCCCACGTCAAGGAGTAT | GAPDH F | GTCAGTGGTGGACCTGACCT |
| Ccl5 R | CCACTTCTTCTCTGGGTTGG | GAPDH R | TCGCTGTTGAACTCAGAGGA |
| Ccl20 F | TTTTGGGATGGAATTGGACAC | | |
| Ccl20 R | TGCAGGTGAAGCCTTCAACC | | |
| Cxcl1 F | TCCAGAGCTTGAAGGTGTTGCC | | |
| Cxcl1 R | AACCAAGGGAGCTTCAGGGTC | | |
| Cxcl2 F | AGTGAACTGCGCTGTCAATG | | |
| Cxcl2 R | TTCAGGGTCAAGGCAAACTT | | |
| Cxcl10 F | GTGAGAATGAGGGCCATAGG | | |
| Cxcl10 R | TTTTTGGCTAAACGCTTTCAT | | |
| Cx3cl1 F | GGCTAAGCCTCAGAGCATTG | | |

| | |
|---|---|
| Cx3cl1 R | CTGTAGTGGAGGGGGACTCA |
| Gcsf F | ATGGCTCAACTTTCTGCCCAG |
| Gcsf R | CTGACAGTGACCAGGGGAAC |
| TNFa F | AAGCCTGTAGCCCACGTCGTA |
| TNFa R | GGCACCACTAGTTGGTTGTCTTTG |
| Tap1 F | TTCCCTCAGGGCTATGACAC |
| Tap1 R | TGGTGGCATCATCCAAGATA |
| Tap2 F | ACAGGACGATGCTGGTGATT |
| Tap2 R | TACCAGGTGGGCGTAGACAT |
| H2d1 F | CATGGTGATCGTTGCTGTTC |
| H2d1 R | CTGGAGCCAGAGCATAGTCC |
| H2k1 F | TCCATCCACTGTCTCCAACA |
| H2k1 R | CTGGAGCCAGAGCATAGTCC |
| B2m F | CTGACCGGCCTGTATGCTAT |
| B2m R | CCGTTCTTCAGCATTTGGAT |
| Erap1 F | CTCATTGGCAGAAACCCAGT |
| Erap2 R | ATGGACGATGAGCCAAGTTC |
| Gapdh F | GCAGTGGCAAAGTGGAGATT |
| Gapdh R | GAATTTGCCGTGAGTGGAGT |
